# Single cell in vivo analysis of type I IFN and NK cell-mediated control of B cell infection densities during acute gammaherpesvirus infection

**DOI:** 10.64898/2026.06.15.732291

**Authors:** Razieh Zargari, Viktoria Rex, Gang Li, J. Craig Forrest, Reinhold Förster, Korbinian Brand, Melanie M. Brinkmann, Stephan Halle

## Abstract

Immunodeficient patients are at risk of severe complications following infection with the human gammaherpesviruses Epstein-Barr virus (EBV) and Kaposi’s sarcoma-associated herpesvirus (KSHV). Due to the strict host specificity of human gammaherpesviruses, murine gammaherpesvirus 68 (MHV68) is widely used as an *in vivo* model to study gammaherpesvirus pathogenesis. While type I interferons (IFNs) and natural killer (NK) cells are known to contribute to antiviral defense during MHV68 infection, how these innate immune mechanisms control infection within lymphoid tissues at the level of individual infected cells remains incompletely understood.

Here, we used an MHV68-DsRed reporter virus to visualize infected cells in the lymph nodes of wildtype and type I IFN receptor-deficient (Ifnar1^-/-^) mice by flow cytometry, immunohistochemistry, and two-photon microscopy. We found that type I IFN signaling plays a major role in limiting both local viral infection in draining lymph nodes and systemic dissemination to the spleen during acute infection. In the absence of IFNAR signaling, infected B cells accumulated to higher numbers, including an increased proportion of infected germinal center phenotype B cells, and this was accompanied by enhanced activation of T and B lymphocytes.

Using two-photon microscopy, we further examined NK cell behavior within infected lymph nodes. NK cells were rapidly recruited to sites of infection but did not form stable clusters or prolonged contacts with infected cells. Nevertheless, NK cell depletion resulted in increased numbers of infected cells, indicating that NK cells contribute to the control of acute MHV68 infection despite the absence of detectable long-lasting interactions with infected cells.

Together, our findings provide a single-cell view of acute gammaherpesvirus infection and innate immune control within lymph nodes *in vivo*. These data refine our understanding of how type I IFN responses and NK cells restrict early gammaherpesvirus spread and shape infection dynamics within lymphoid tissues.

**Author Summary:** Certain viruses can remain in the body for life and cause little harm in healthy individuals. However, when the immune system is weakened, these infections can lead to cancer or other life-threatening diseases. To study human gammaherpesviruses within the living body, scientists often use a closely related mouse virus to understand how the immune system controls these infections.

In this study, we tracked virus-infected cells within lymph nodes, important sites of the immune responses. Using a fluorescent virus together with advanced imaging techniques, we visualized infected cells directly in intact tissues. We found that type I interferons, a key component of the early antiviral defense, strongly limited the number of infected cells. Mice that could not respond to type I interferons developed substantially larger infections.

We also investigated the role of natural killer cells, immune cells that provide rapid protection against viral infections. Although these cells quickly accumulated at sites of infection, they rarely formed prolonged contacts with infected cells.

Together, our findings provide a detailed view of how innate immune defenses restrict gammaherpesvirus infection within lymph nodes. By revealing how early immune responses limit viral spread, this work improves our understanding of mechanisms that protect against severe herpesvirus-associated disease.

## Introduction

Gammaherpesviruses are oncogenic viruses that infect many different cell types, including epithelial cells, macrophages, and dendritic cells. B lymphocytes are thought to represent the major target population. Epstein-Barr virus (EBV) and Kaposi’s sarcoma-associated herpesvirus (KSHV) are two human gammaherpesviruses that establish chronic infections in their hosts. While KSHV is primarily endemic in parts of Africa, Asia, and South America, approximately 95% of adults worldwide are latently infected with EBV [1,2]. EBV infection during adolescence or adulthood can lead to the development of infectious mononucleosis, and is associated with the development of several autoimmune diseases, including multiple sclerosis [3].

In immunocompromised individuals, EBV infection can cause severe complications, including B-cell malignancies like post-transplant lymphoproliferative disease [4]. In rare cases, individuals are unable to control EBV infection and reactivation, which leads to the development of chronic active EBV (CAEBV) disease. During CAEBV disease, EBV^+^ lymphocytes infiltrate multiple organs, leading to elevated viral load in the peripheral blood. To date, hematopoietic stem cell transplantation remains the only definitive therapy [5,6]. Another risk for severely immunodeficient patients is gammaherpesvirus-triggered secondary hemophagocytic lymphohistiocytosis (HLH) [7]. KSHV, on the other hand, is associated with the development of Kaposi’s sarcoma, and the lymphoproliferative disorders multicentric Castleman disease (MCD) and primary effusion lymphoma [8,9]. Hence, it is important to improve our understanding of how an intact or attenuated immune response affects gammaherpesvirus replication, spread, and host control mechanisms.

Many aspects of human gammaherpesvirus pathogenesis cannot be studied in the natural human host due to the strict host-specificity of these viruses, and we therefore depend on suitable model systems. Murid herpesvirus 4 (MuHV-4), commonly referred to as murine gammaherpesvirus strain 68 (MHV68), is a natural pathogen of mice and has been widely used to study gammaherpesvirus infection *in vivo [10–14].* Similar to human gammaherpesviruses, MHV68 infects and exploits B cells to establish latency and has similarly developed strategies to evade innate and adaptive immune responses to promote lytic infection [15–18].

The type I interferon (IFN) system and natural killer (NK) cells are two major players during the innate immune response. Type I IFNs play a fundamental role in protecting the host after viral infection, as demonstrated by many studies in mice lacking the type I IFN receptor (IFNAR) [19]. Sensing of viral infection by pattern recognition receptors (PRRs) induces the secretion of type I IFNs and subsequent activation of type I IFN receptor (IFNAR) signaling and transcription of hundreds of interferon-stimulated genes (ISGs) with diverse functions [20]. Further, the type I IFN response also supports the initiation and shaping of the innate and adaptive immune responses after gammaherpesviral infection by directly or indirectly affecting dendritic cells, B and T lymphocytes, and NK cells [21]. Following activation, NK cells contribute to the antiviral immune response through distinct effector functions [22]. NK cells primarily eliminate infected cells through contact-dependent cytotoxicity via immunological synapse formation [23], death receptor signaling [24], and the release of granzymes and perforin [25,26], while also secreting pro-inflammatory cytokines and chemokines that activate other immune cells [27]. The importance of NK cell-mediated control of EBV infection is highlighted in studies of the immune response [28] and is further characterized using EBV infected human tonsils and PBMCs [29–31].

Previous studies in *Ifnar1*^-/-^ mice have shown that the type I IFN response is crucial for controlling acute MHV68 infection *in vivo* [32–34]. Notably, vaccination with a replication-deficient MHV68 protects mice lacking type I IFN responses from severe disease [35]. While several studies have investigated MHV68 tropism, spread, and the contribution of innate and adaptive immunity [36–41], the immune response in *Ifnar1^-/-^* mice has not yet been investigated at the single cell level. Consequently, kinetics of viral dissemination, mechanisms of acute infection control, and the role of NK cells in restraining MHV68 in lymph nodes remain unresolved.

Here, we demonstrate the crucial role of the type I IFN and NK cell response *in vivo* during acute MHV68 infection on the single cell level by combining two-photon microscopy, flow cytometry, and immunohistochemistry of infected lymph nodes.

## Results

### MHV68 infects the popliteal and para-aortic lymph nodes after subcutaneous footpad injection

In this study, we used the recombinant MHV68-DsRed virus strain, which expresses the red fluorescent protein DsRed, to visualize MHV68-infected cells following in vivo footpad injection (Fig 1). Subcutaneous (s.c.) footpad injection is a well-established model in mice to study the local immune response in draining lymph nodes following infection and has previously been used to study MHV68 infection [42]. In this model, the virus particles are injected subcutaneously into the footpad of anesthetized mice. Viral particles reach the draining lymph node of the footpad and popliteal lymph node via lymph drainage and migratory DCs, leading to the generation of local infection and an immune response. The local infection in the popliteal lymph nodes can then reach downstream draining lymph nodes, including the para-aortic lymph nodes, and eventually lead to systemic spread by entering blood circulation via the *vena cava* after spreading through the lymph node chain (Fig 1) [43,44].

**Fig 1.**
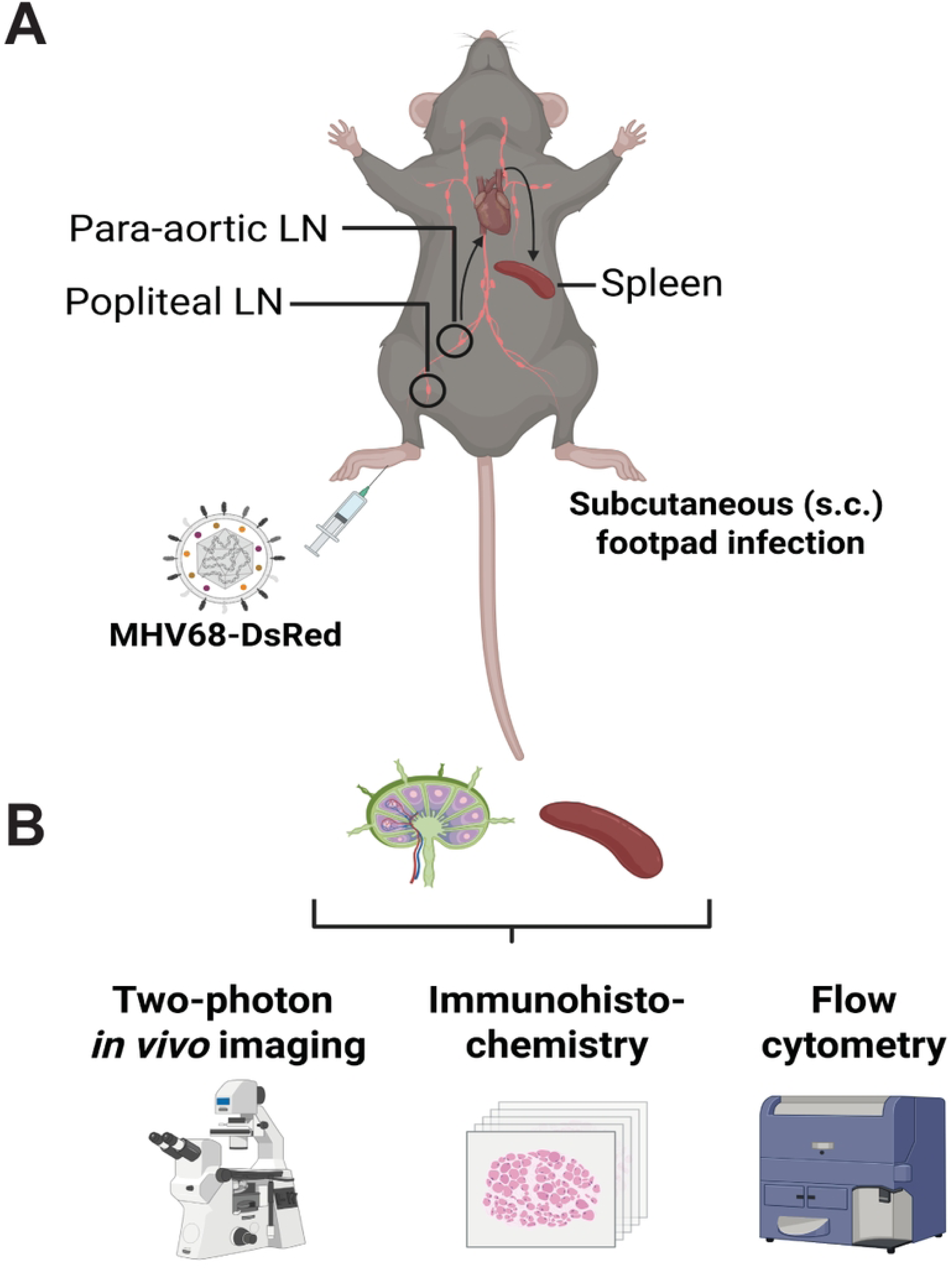
Visualization of MHV68 infection in mice. Scheme of the experimental setup. **(A)** Mice were infected subcutaneously (s.c.) in the footpad with an MHV68-DsRed reporter virus and sacrificed on day 1, 3, 5, or 7 post-infection for extraction of the draining and non-draining lymph nodes and the spleen. Pink lines show MHV68-DsRed trafficking following s.c. infection. Shown are the popliteal and para-aortic lymph nodes and the spleen. **(B)** Organs are analyzed by two-photon microscopy, immunohistochemistry, and flow cytometry. Created in BioRender. https://BioRender.com/iw60fy9

Following s.c. footpad infection with MHV68-DsRed, we searched for DsRed^+^ virus-infected cells in the popliteal and para-aortic lymph nodes by two-photon microscopy and flow cytometry (Fig 2A). On day 5 p.i., we could clearly identify a population of bright DsRed^+^ virus-infected cells, with around 0.03% of total live single cells expressing high levels of DsRed in popliteal lymph nodes (Fig 2B). DsRed was expressed with different mean fluorescence intensities per cell, and no clear DsRed expression was seen on day 1 and 2 post infection (data not shown). DsRed^+^ cells were also detectable in para-aortic lymph nodes at day 5 and 7 p.i., indicating the spread of infection to downstream lymph nodes (Fig 2C).

**Fig 2.**
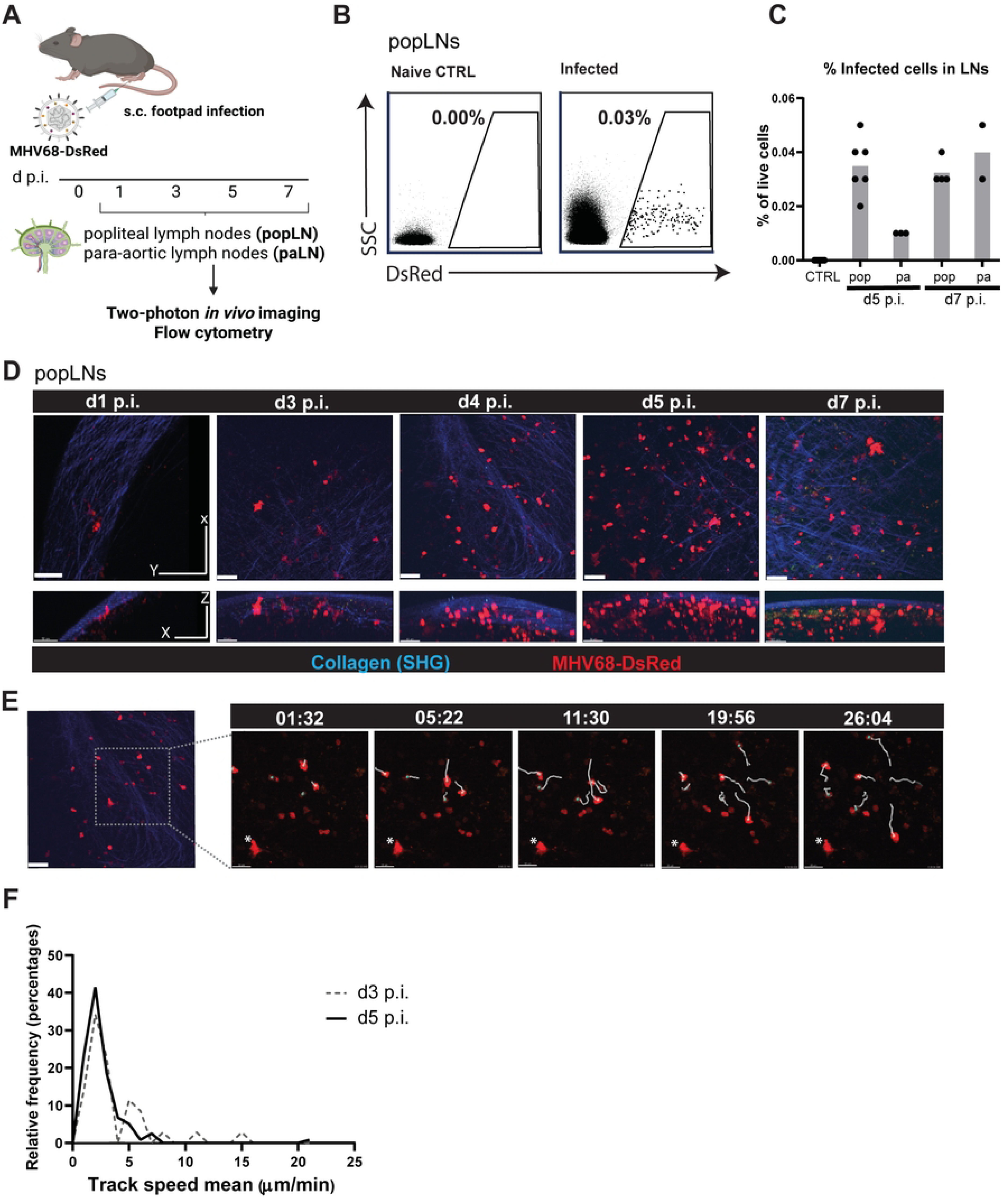
Flow cytometry and two-photon microscopy analysis of MHV68-DsRed-infected cells in lymph nodes. **(A)** Wildtype mice were infected subcutaneously (s.c.) in the footpad with an MHV68-DsRed reporter virus (day 0) and sacrificed on day 1, 3, 5, or 7 post-infection for extraction of the lymph nodes (LNs) followed by two-photon imaging and flow cytometry. Created in BioRender. https://BioRender.com/lf7f55j **(B)** Identification of MHV68-DsRed-infected cells in LNs. Representative flow cytometry plots showing the DsRed^+^ cell population in popLNs from naive control (CTRL) (left) and MHV68-DsRed-infected (right) mice 5 d.p.i. gated on live single cells. **(C)** Percentage of MHV68-DsRed-infected cells in popliteal (pop) and para-aortic (pa) LNs on days 5 and 7 p.i. Each dot represents a popliteal LN or pooled para-aortic LNs for each mouse. **(D)** Representative two-photon microscopy images of infected popliteal LNs at the indicated time points post-infection; upper row shows the top view and the lower row the lateral view for each image. The second-harmonic generation (SHG) indicates collagen. Scale bar: 50μm. **(E)** Representative two-photon images showing the tracking of motile (indicated with white lines) and stationary (indicated with *) infected cells in the popliteal LN over time. **(F)** Analysis of track speed of MHV68-DsRed-infected cells in the popliteal LNs at days 3 and 5 p.i based on single cell tracks. N = 1-3 mice per time point pooled from 3 independent experiments.

### Two-photon microscopy of the popliteal lymph node identifies morphologically distinct MHV68-infected cell types

Next, we studied the motility characteristics of gammaherpesvirus-infected cells in vivo in lymph nodes. Although it has been presumed that gammaherpesvirus-infected lymphocytes are motile and could potentially facilitate the spread of infection by migrating to distal organs, their motility has never been observed in living virus-infected tissues. Two-photon microscopy analysis of infected cell motility in the popliteal lymph nodes at different time points post infection revealed the presence of two morphologically distinct infected cell types: stationary and motile cells (Fig 2D-F). The stationary DsRed+ MHV68-infected cells were first detected at day 1-2 p.i., with a morphology and positioning similar to macrophages (Fig 2D). The stationary cells were attached to the lymph node capsule, corresponding to CD169^+^ sub-capsular sinus (SCS) macrophages observed in histological analysis (data not shown). On the other hand, the motile infected cells were detected after day 3 p.i. and peaked at day 5 p.i. in the popliteal lymph nodes. Furthermore, the majority of infected cells at day 5 p.i. were motile. Nevertheless, we observed that motile infected cells exhibited predominantly slow, confined motility patterns (Fig 2F; S1 Movie 1). Notably, two-photon microscopy of the infection site allowed clear identification of even the very few initially infected cells on days 1-3 post infection.

### The type I interferon response regulates gammaherpesvirus infection of the lymph nodes in vivo

MHV68 antagonizes the type I IFN response in various ways in order to promote lytic infection, emphasizing the evolutionary importance of this antiviral pathway [45–48]. Nevertheless, the type I IFN response plays a crucial role in controlling MHV68 infection in mice. Previous *in vivo* studies revealed that *Ifnar1^-/-^* mice are highly susceptible to MHV68 infection. *Ifnar1^-/-^* mice have a dysfunctional type I IFN receptor and cannot respond to secreted type I IFN, which makes them highly susceptible to viral infection. MHV68 infection spreads faster in *Ifnar1^-/-^*mice, and the mice ultimately succumb due to their inability to control the acute infection [38,49]. However, the number of virus-infected lymphocytes and the impact of type I IFN deficiency on the development of the adaptive immune response during acute MHV68 infection remain unclear [21]. Thus, we analyzed the role of the type I IFN response in controlling acute MHV68 infection in the lymph nodes at the single cell level. Additionally, we sought to compare the adaptive immune response following MHV68 infection in WT and *Ifnar1^-/-^* mice. Towards this aim, WT and *Ifnar1^-/-^* mice were infected with MHV68-DsRed via s.c. footpad infection. We then analyzed the local infection and immune response in popliteal lymph nodes and systemic spread to the spleen.

On day 1 p.i., histological analysis of popliteal lymph nodes revealed virus-infected cells in the subcapsular sinus region in both WT and *Ifnar1^-/-^* mice (data not shown). After 3 and 5 days p.i., popliteal lymph nodes from *Ifnar1^-/-^* mice had substantially more infected cells than those from WT mice (Fig 3A). This directly demonstrates that IFNAR regulates the density of infected B cells *in vivo*. Additionally, we observed accumulation of CD11b^+^ cells around the B cell follicles and in the interfollicular regions of the infected lymph nodes, with denser CD11b+ cell infiltration in the *Ifnar1^-/-^* popliteal lymph nodes (Fig 3B). CD11b (integrin alpha M) is expressed on the surface of many leukocytes, including monocytes, dendritic cells, and NK cells, and is involved in adhesion, migration, and accumulation of leukocytes in the inflammatory response [50]. These data indicate that MHV68 initially infects the same target cells in WT and *Ifnar1^-/-^*popliteal lymph nodes, but viral amplification on the single cell level *in vivo* is more efficient in the absence of type I IFN signaling.

**Fig 3.**
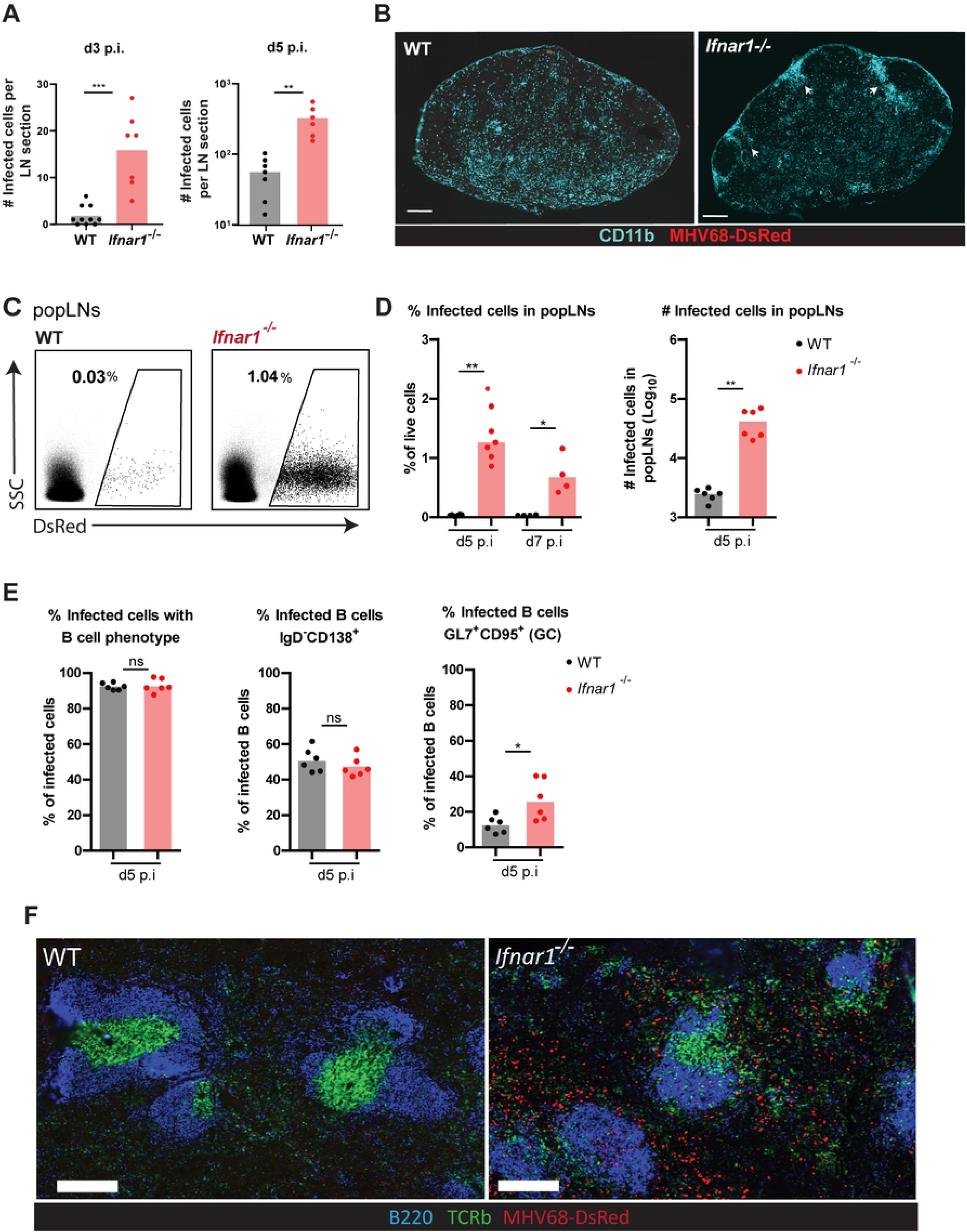
Role of the type I IFN response in controlling acute MHV68 infection in lymph nodes and spleen. WT and *Ifnar1^-/-^* mice were infected with MHV68-DsRed via s.c. footpad infection. The popliteal LNs were isolated at the indicated timepoints p.i. for flow cytometry and immunohistochemistry (IHC). **(A)** Automated counting analysis of the number of infected cells in popliteal LN sections from WT and *Ifnar1^-/-^* mice at days 3 and 5 p.i. by IHC. Each dot represents a LN section. **(B)** Representative IHC images of popliteal LN sections from WT and *Ifnar1^-/-^* mice at day 3 p.i. labeled with an antibody specific for CD11b. Arrows indicate areas with an accumulation of CD11b^+^ cells. Scale bar: 200 μm. **(C)** Representative flow cytometry plots showing the DsRed^+^ population in popliteal LNs of WT and *Ifnar1^-/-^*mice on day 5 p.i. gated on live single cells. **(D)** The frequency of DsRed^+^ (infected) cells in popliteal LNs in WT and *Ifnar1^-/-^* mice on days 5 and 7 p.i. (left panel) and the total numbers of DsRed^+^ infected cells in popliteal LNs in WT and *Ifnar1^-/-^* mice on day 5 p.i. (right panel). Each dot represents an individual LN. **(E)** Analysis of the phenotype of infected cells in WT and *Ifnar1^-/-^* mice in popliteal LNs at day 5 p.i. by flow cytometry. The surface expression of CD19, IgD, CD138, GL7, and CD95 was used as markers to identify B cells with plasmablast or germinal center phenotype as indicated. Each dot represents a LN. **(F)** The Fig shows representative images of spleens from WT and *Ifnar1^-/-^* mice following MHV68-DsRed s.c. footpad infection. Spleens were isolated 5 d.p.i., fixed in PFA and analyzed by IHC. The sections were labeled with anti-B220 and anti-Tcrβ antibodies to visualize B cell (blue) and T cell (green) zones, respectively. Scale bar: 100 μm. N = 3 mice per time point per condition. **A, D, E** N = 2-3 mice per time point per condition for flow cytometry and IHC analysis. The bars indicate the mean. Statistical analysis was performed using the Mann-Whitney test. ns, not significant; p-values <0.05 (*), <0.005(**), <0.0005(***).

By flow cytometry, we also found that the frequency and total number of DsRed+ infected cells were higher in Ifnar1-/- mice than in WT mice at days 5 and 7 p.i. (Fig 3C-D), indicating more severe acute infection as well as possible inflammation in the absence of type I IFN signaling. Interestingly, analysis of infected cell phenotypes by flow cytometry revealed that the majority (∼90%) of the infected cells in WT and *Ifnar1^-/-^* conditions were CD19^+^ B cells, including IgD^-^CD138^+^ plasmablasts and GL7^+^CD95^+^ germinal center (GC) B cells. While there was no significant difference between the frequencies of infected plasmablast cells in *Ifnar1^-/-^* or WT mice, *Ifnar1^-/-^* mice showed higher frequencies of infected B cells with a GC phenotype (Fig 3E). In summary, we observed that in the absence of type I IFN signaling, B cells were still the main target population, and MHV68 cellular tropism did not change significantly.

### The type I IFN response protects the host from uncontrolled systemic spread

Next, we investigated the systemic spread of acute MHV68 infection in the absence of the type I IFN response from the primary infected lymph nodes to the spleen (Fig 3F). Infection could spread systemically from the primary site of infection to the spleen, potentially either via the dissemination (transmission) of free virus particles or via the migration of infected cells from the primary site. Histological analysis of the spleen revealed that MHV68-DsRed-infected cells were detectable in the spleen at day 5 post s.c. infection in *Ifnar1^-/-^* mice (Fig 3F) with extensive infection at day 7 p.i. (data not shown). We could not detect DsRed^+^-infected cells in splenic sections of WT mice at any analyzed time points (data not shown). The infected cells in *Ifnar1^-/-^* spleens were distributed throughout, including the white pulp and red pulp, indicating the importance of the type I IFN response in containing acute MHV68 infection and preventing extensive systemic infection. Taken together, the MHV68-DsRed *in vivo* infection model clearly shows type I IFN-mediated protection against massive viral spread at the single cell level, as observed by two-photon microscopy and flow cytometry.

### MHV68-induced regulation of interferon-stimulated genes in vivo

Interferon-triggered antiviral resistance mechanisms prevent virus spread. Various interferon-stimulated genes (ISG) are expressed in response to type I IFN and prevent viral replication and cell-cell spread. Although the majority of cells are able to respond to type I IFN via their IFNAR receptors, the cellular response to IFN is not homogeneous and different cell types in tissues can respond differentially to secreted type I IFN. How ISG dynamics in lymph nodes are regulated by IFNAR signaling *in vivo* has previously not been shown. Thus, we were interested to understand the response of the total lymph node cell population as well as B cells to type I IFN upon MHV68 infection.

To investigate the role of type I IFN-dependent ISG expression in protecting mice from MHV68-DsRed acute infection, we analyzed the expression of the ISGs *ISG15, MX1, CXCL10*, and *mOASL2* in the total isolated cell population of infected lymph nodes from WT and *Ifnar1^-/-^*mice at 6 and 18-24 hrs post-infection (Fig 4). Expression of *ISG15, MX1, CXCL10,* and *mOASL2* was very low 6 hrs p.i., but increased in lymph nodes of WT mice 18-24 hrs p.i. (Fig 4A). Interestingly, the expression of *ISG15* and *MX1* was very low in *Ifnar1^-/-^* mice at 18-24 hrs p.i., while *CXCL10* and *mOASL2* expression was detectable. We could not detect *IFNα* and *IFNβ* gene expression at any of the analyzed time-points (data not shown). These data fit to our observations on the single cell level, where the initial number of virus-infected cells was similar in WT and *Ifnar1^-/-^* mice on day 1 after infection. Because induction of the type I IFN response and subsequent ISG expression requires time, the phenotype of WT and *Ifnar*1^-/-^mice thus appears similar regarding IFNα and IFNβ gene expression very early after infection.

**Fig 4.**
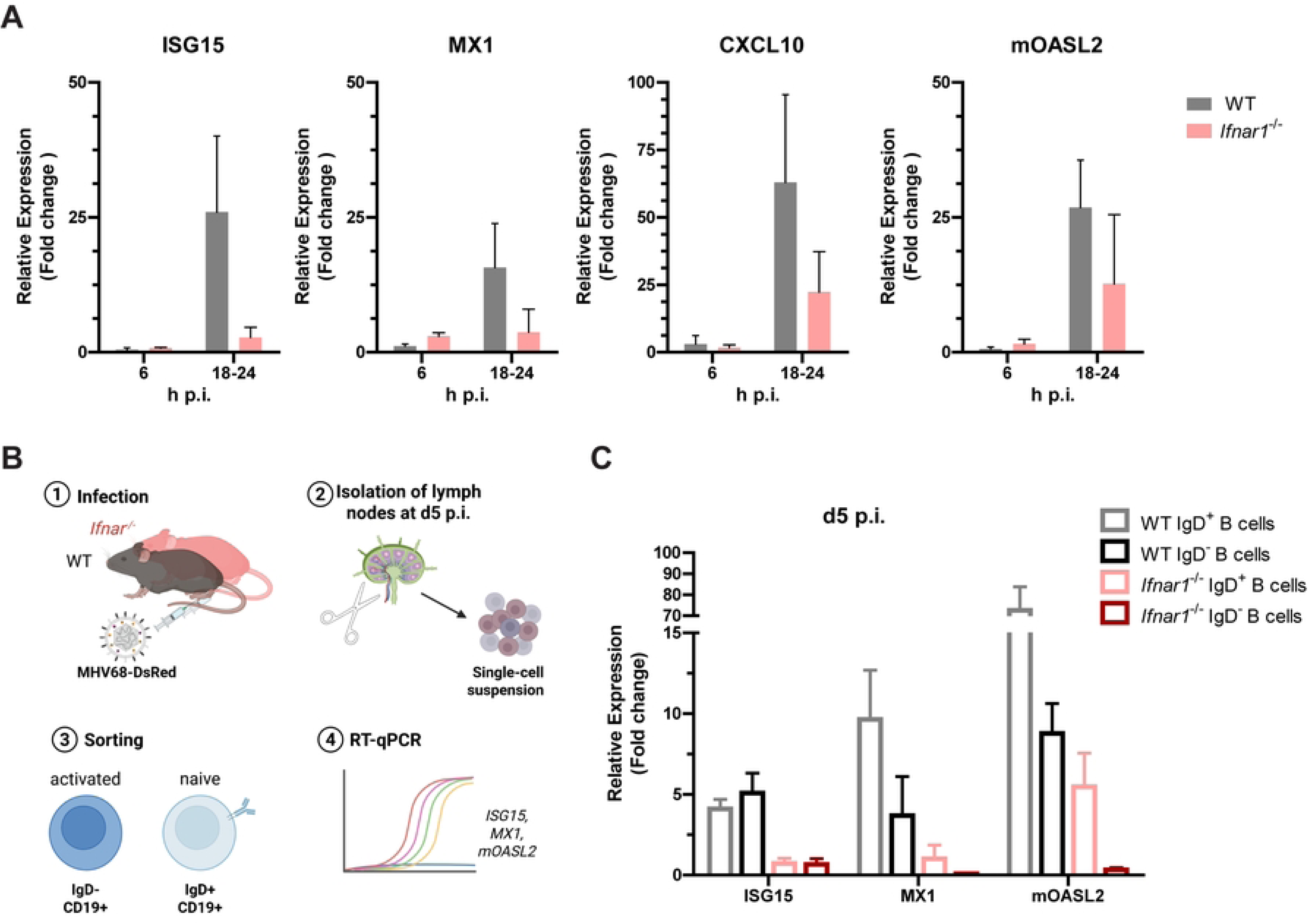
Expression of interferon-stimulated genes (ISGs) in MHV68-DsRed-infected LNs. **(A)** Wildtype and *Ifnar1^-/-^* mice were infected with MHV68-DsRed via s.c. footpad infection and popliteal LNs and para-aortic LNs were isolated. Total RNA was isolated from pooled lymph node single-cell suspensions, cDNA generated and expression of the indicated ISGs was analyzed by RT-qPCR. Fold expression over non-infected samples was calculated by analysis of relative expression compared to naive control for each condition. Data were first normalized to GAPDH expression in all conditions. Created in BioRender. https://BioRender.com/uircjcp **(B)** WT and *Ifnar1^-/-^* mice were infected with MHV68-DsRed via s.c. footpad infection and LNs were isolated, followed by sorting of the CD19^+^IgD^-^(activated B cells) and CD19^+^IgD^+^ (naive B cells) cells and RT-qPCR analysis of the indicated ISGs. **(C)** Relative expression of ISGs in B cells of WT and *Ifnar1^-/-^* mice 5 days p.i.. **A, C** Data were pooled from 3 independent experiments using n=4 mice per condition per time point. Mean + SD are shown. No statistical tests were planned.

We were further interested in the level of ISG expression in activated and naive B cells in the lymph nodes of WT and *Ifnar1^-/-^* mice at the peak of infection on day 5. We used high versus low IgD surface expression to differentiate between naive and activated B cells (Fig 4B). Notably and similarly to what we had observed in lymph nodes at 18-24 hrs p.i., we detected increased expression of *ISG15*, *MX1*, and *mOASL2* in sorted B cells isolated from the lymph nodes of WT mice, while the expression in B cells from *Ifnar1^-/-^* mice was decreased (Fig 4C). Interestingly, naive B cells exhibited substantially higher *mOASL2* expression than activated B cells in both WT and *Ifnar^-/-^* mice, although the overall magnitude of this fold change was reduced, as previously described. A similar pattern was observed for *MX1*, but not *ISG15*.

In summary, we observed that expression of *MX1* and *ISG15* was decreased in *Ifnar1^-/-^* vs WT mice, not only in the total lymph node populations at early time points p.i., but also in isolated B cells at the peak of infection, showing that during MHV68 infection *in vivo*, naive and activated B cells are responsive to type I IFN in an IFNAR-dependent manner.

### Analysis of the adaptive immune response following MHV68-DsRed infection in WT and *Ifnar1^-/-^* mice

The type I IFN response also protects the host from viral infections by contributing to the initiation and shaping of an effective adaptive immune response. Thus, we next investigated the development of the adaptive immune response following MHV68 infection in the presence or absence of type I IFN signaling. To analyze the adaptive immune response in single lymph nodes of WT and *Ifnar1^-/-^* mice following MHV68-DsRed infection, a multicolor spectral flow cytometry panel was designed to identify several key populations of cell-mediated and humoral immune responses, including effector CD8^+^ T cells, follicular helper CD4^+^ T cells (Tfh), and germinal center (GC) B cells.

In EBV-triggered secondary HLH, massive CD8^+^ T cell activation is observed, and patients have a very dismal prognosis. Thus, we aimed to better understand T cell and B cell activation in acute MHV68 infection (Fig 5 A-C). We analyzed the CD8^+^ T cell population in the popliteal lymph nodes of WT and *Ifnar1^-/-^* mice at different time points and used CD44 and CD62L markers to identify effector CD8^+^ T cells as the main effector population in the antiviral cell-mediated adaptive response (Fig 5D). Surprisingly, effector CD8^+^ T cells were developed and detectable in both conditions starting at day 5 p.i., showing induction of T cell immunity early after infection even in the absence of type I IFN signaling. Notably, we observed that the frequency of total CD8^+^ T cells was significantly higher in *Ifnar1^-/-^* mice at days 3 and 5 p.i. compared to WT mice. Interestingly, the frequency of the naive CD8^+^ T cells population was significantly higher in WT compared to *Ifnar1^-/-^* mice at days 5 and 7 p.i., while the frequency of CD62L^-^CD44^+^ effector CD8^+^ T cells was remarkably higher in *Ifnar1^-/-^* mice compared to WT (at days 5 and 7 p.i) (Fig 5D). These data suggest that in the absence of a type I IFN response, other inflammatory pathways could play a compensatory role in T cell activation and priming.

**Fig 5.**
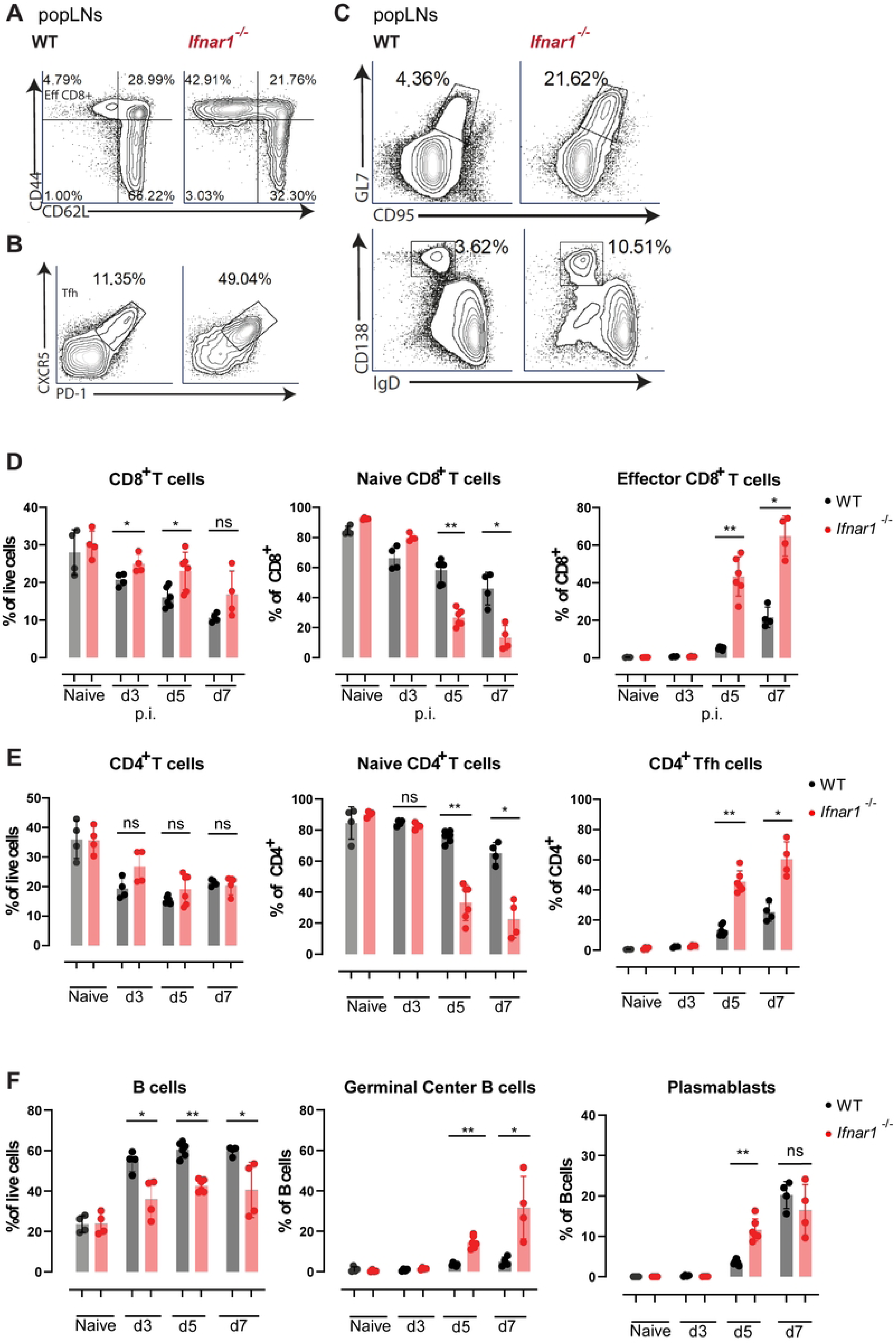
Analysis of the humoral immune response in popliteal LNs of WT and *Ifnar1^-/-^* mice during acute MHV68-DsRed infection. Mice were infected with MHV68-DsRed and the popliteal LNs were isolated and analyzed by flow cytometry. **(A)** Representative plots showing the naive CD62L^+^ CD44^-^and effector CD62L^-^ CD44^+^ CD8^+^ T cell populations in popliteal LNs on day 5 p.i. **(B)** Representative plots showing the PD-1^+^CXCR5^+^ Tfh population in WT and *Ifnar1^-/-^* mice at day 5 p.i.. **(C)** Representative plots showing the GL7^+^CD95^+^ germinal center B cells (upper row) and the IgD^-^CD138^+^ plasmablasts (bottom row) in the popliteal LNs of WT and *Ifnar1^-/-^* mice on day 5 p.i.. **(D-F)** Frequencies of **(D)** total, naive, and effector CD8^+^ T cells, **(E)** total, naive, and Tfh CD4^+^ T cells and **(F)** total B cells, germinal center B cells, and plasmablasts in WT and *Ifnar1^-/-^* popliteal LNs at day 3, 5, and 7 p.i. analyzed by flow cytometry. Each dot represents a LN. N = 2-3 mice analyzed per time point per condition in 3 independent experiments. Mean +- SD are shown. Statistical analysis was performed using the Mann-Whitney test. ns, not significant; p-values <0.05 (*), <0.005(**).

Next, we were interested in studying the development of the B cell immune response in the presence or absence of type I IFN signaling. Germinal center B cells and Tfh cells, as well as plasmablasts, are the main effector populations of the humoral immune response, which leads to the production of high-affinity antibodies. To understand their contribution during MHV68 infection, we analyzed the populations of Tfh, GC B cells, and plasmablasts in the popliteal lymph nodes of WT and *Ifnar1^-/-^* mice (Fig 5E-F). Starting from day 5 p.i., we observed that both CD4^+^ Tfh cells as well as the GC B cells developed in WT and KO mice. Interestingly, the frequency of the Tfh population, as well as GC B cells, was significantly higher in *Ifnar1^-/-^* mice compared to WT at days 5 and 7 p.i., indicating the development of a more robust T cell-dependent B cell activation in *Ifnar1^-/-^* mice at the peak of infection. In addition, we observed that *Ifnar1^-/-^* had significantly higher frequencies of plasmablasts on day 5 p.i. compared to WT mice.

Altogether, we observed that the effector populations of the cell-mediated adaptive immune response massively expanded following MHV68 infection in the popliteal lymph nodes of WT and *Ifnar1^-/-^* mice *in vivo*. Although type I IFN plays an important role in initiating the adaptive immune response, the presence of more MHV68-infected cells in the *Ifnar1^-/-^* mice, as well as other type I IFN independent inflammatory mechanisms, could both have supported a strong antiviral immune cell response in the absence of IFN-I signaling. Notably, the very strong CD8^+^ T cells response in IFNAR-deficient mice might provide a good model system to study secondary gammaherpesvirus-induced HLH.

### The role of NK cells in controlling acute MHV68 infection

It has been shown that NK cells expand and become activated following MHV68 infection *in vivo*, but their depletion results in no significant difference in viral loads between depleted and control groups in mice [51,52]. Furthermore, the role of NK cells in restricting MHV68 in lymph nodes was previously demonstrated by Lawler et al., who showed that NK cells are recruited to infected popliteal lymph nodes and that their depletion increases the number of lytically infected subcapsular sinus macrophages [38]. In the present study, using two-photon in vivo microscopy, we aimed to visualize NK cell-MHV68-infected cell interactions in greater detail. We infected Ncr1^gfp^ reporter mice with MHV68-DsRed virus via s.c. footpad infection, and analyzed the popliteal lymph nodes for up to 5 days p.i. (Fig 6A-D). The analysis of the popliteal lymph nodes in Ncr1^gfp^ mice with two-photon microscopy showed that the GFP^+^ NK cells were recruited to the site of infection starting at day 1 p.i. (Fig 6A-B), consistent with the findings of Lawler et al. Interestingly, our analysis of NK cell motility in intact lymph nodes demonstrated that the majority of NK cells were motile, particularly on days 1 and 3 post-infection (Fig 6C). Nevertheless, NK cells formed no clusters or long-lasting contacts with stationary or motile infected cells, and we did not observe any NK cell-mediated contact-dependent killing of infected cells throughout the analysis at these time points (S2 Movie 2).

**Fig 6.**
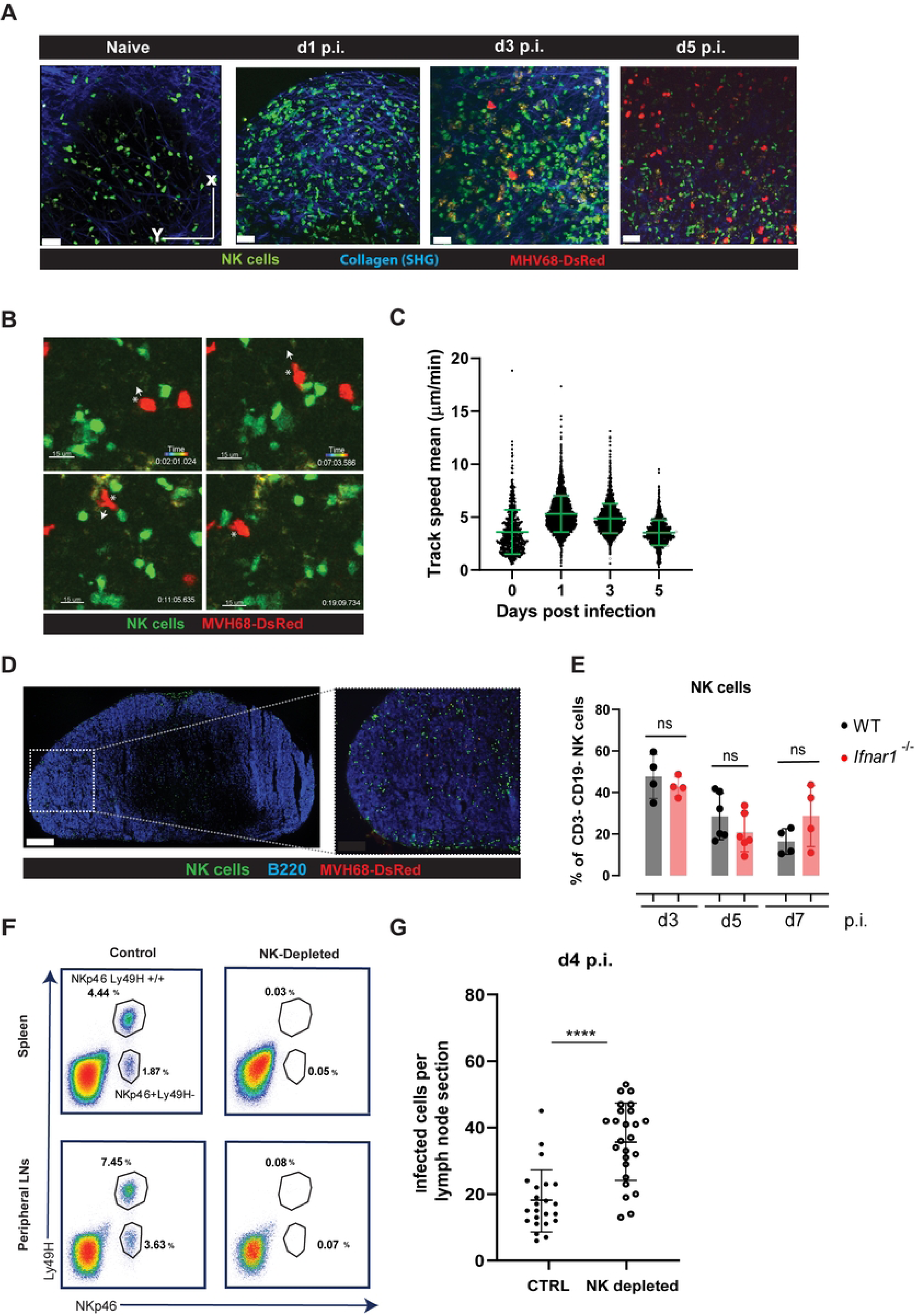
Analysis of Natural Killer (NK) cell recruitment to and interaction with MHV68-infected cells. **(A-D)** Ncr1^gfp^ reporter mice, allowing fluorescent visualization of NK cells, were infected with MHV68-DsRed s.c. via the footpad. The popliteal LNs were isolated on day 1, 3, and 5 p.i. and analyzed by two-photon microscopy. **(A)** The representative images of popliteal LNs show the NK cells (green) in the infected (red) popliteal LNs at different time points p.i.. The second-harmonic generation (SHG) indicates collagen (blue). Scale bar: 50 μm. **(B)** The representative images show the motility and interactions between NK cells and motile MHV68-DsRed-infected cells. The arrow indicates the direction of the motile MHV68-DsRed-infected cell. Scale bar: 30 μm. **(C)** Analysis of NK cell motility in the popliteal LNs. Each dot represents a track. 2-3 movies per LN were analyzed. **(D)** Representative image of MHV68-DsRed-infected popliteal LN on day 3 in Ncr1^gfp^ mice analyzed by IHC. The sections were labeled with anti-B220 to visualize B cell zones (blue). Scale bar: 200 μm. **(E)** Frequencies of CD3^-^CD19^-^ NK cells in popliteal LNs on days 3, 5, and 7 p.i. in WT and *Ifnar1^-/-^* (KO) mice. Each dot represents a LN. **(F-G)** WT mice were infected with MHV68-DsRed via s.c. footpad infection. NK cells were depleted by i.p. injection with an anti NK1.1 antibody (250 µg) 1 day prior to infection, and at 0, 2, and 4 days p.i. NK depletion was confirmed by flow cytometry analysis of spleen and peripheral LNs in control and NK cell-depleted mice for each experiment. **(F)** Representative plots showing the NK cell gate (CD19 CD3^-/-^ population) identified by the NKp46 marker in the spleen and pooled peripheral LNs of NK-depleted and control mice on day 4 p.i.. **(G)** IHC analysis of the number of MHV68-infected cells in the popliteal LN sections on day 4 p.i. in control and NK-depleted mice. Each dot represents a LN section. Average CTRL= 17.96± 9,36, NK depleted= 35.91±12.14. Cohen’s d=1.657 (large effect). **C, E, G** The error bar shows the mean ± SD. 3 independent experiments performed with **C** n=3 mice, **E** n=2-3 mice per time point per condition and **G** n=4 mice per group. Statistical analysis was performed using the Mann-Whitney test. ns, not significant, p <0.0001 (****).

The analysis of the recruitment and localization of NK cells following MHV68-DsRed infection in Ncr1^gfp^ mice revealed that NK cells were recruited and distributed in different regions of the popliteal lymph nodes including the B-cell zones (Fig 6D). In addition, by flow cytometry, we did not observe any significant differences in the percentages of NK cells between WT and *Ifnar1^-/-^* mice at days 3, 5, and 7 p.i (Fig 6E), showing that recruitment of NK cells to lymph nodes following MHV68 infection is not remarkably affected in the absence of the type I IFN response.

By closely observing individual NK cells over time using two-photon microscopy, we did not observe any obvious NK cell-mediated contact-dependent killing of infected cells (data not shown). Nevertheless, NK cells could still play a role to restrict MHV68 infection in a contact-independent manner via secretion of inflammatory cytokines. To address this possibility, we studied potential contact-independent effector functions of NK cells during MHV68 early acute infection, before the development and expansion of antigen-specific effector CD8^+^ T cells. To this aim, we depleted the NK cells by injecting anti-NK1.1 antibody in WT C57BL/6 mice and analyzed the number of infected cells on day 4 p.i. (Fig 6F). Consistent with Lawler et al. we observed that the number of infected cells in the popliteal lymph nodes was significantly higher in NK-depleted mice compared to the control group (Fig 6G).

In summary, we did not observe any NK-cell contact-mediated killing or long-lasting interactions between NK cells and MHV68-infected cells in C57BL/6 mice. Our findings are consistent with prior reports revealing that NK cells contribute to the control of acute MHV68 infection in the popliteal lymph nodes *in vivo*. However, the underlying mechanism remained unclear. By directly visualizing NK cell behavior in real time, we demonstrate that NK cells are actively recruited and remain motile within the lymph node, including B cell zones, and that they do not form stable, long-lasting contacts with MHV68-infected cells suggesting that NK cells control MHV68 infection by mechanisms independent of long-lasting NK-cell-target cell interactions.

## Discussion

Uncontrolled gammaherpesvirus replication and spread can trigger excessive T cell activation and life-threatening cytokine storms in human patients. Therapeutic suppression of immune cell activity in such emergencies, however, risks impairing antiviral immunity, presenting a challenging clinical dilemma [53]. To better understand this complex virus-host interaction, we investigated the role of the innate immune response in controlling acute gammaherpesvirus infection in more detail by taking advantage of the murine MHV68 infection model. Gammaherpesviruses are oncogenic, lymphotropic viruses. They infect B-lymphocytes and take advantage of B cell physiology and activation pathways to establish a latent reservoir in memory B cells. In establishing and maintaining this reservoir, gammaherpesviruses require a fine balance with the host immune system.

In this study, we first investigated the type I and NK cell response in controlling acute MHV68 infection in the lymph nodes at the single cell level. By two-photon microscopy and flow cytometry, we could visualize cells infected by MHV68-DsRed reporter viruses in WT and transgenic mice. We observed that in the absence of the type I IFN response, the number of MHV68-infected B cells with a germinal center (GC B cell) phenotype (CD19+, IgD-, GL7+, and CD95+) was increased. Additionally, on day 5 post-infection in the lymph nodes, the frequencies of effector cell populations of adaptive cell-mediated (effector CD8+ T cells) and the humoral immune response (GC B cells, CD4+ follicular helper (Tfh) cells) were significantly higher in *Ifnar1-/-* mice. We detected ISG expression in infected lymph nodes early after infection and in B cells of WT mice at the peak of infection, whereas expression was markedly decreased in *Ifnar1-/-* mice. This indicated that type I IFN-dependent expression of ISGs may protect B cells at the peak of infection and help control the acute infection.

The type I IFN response plays a critical role in protecting individual cells from virus replication, as well as initiating inflammation and shaping an effective antiviral immune response. Type I IFN deficiencies can include deficiencies in type I IFN receptors (IFNAR1, IFNAR2) as well as signaling mediators downstream the IFNAR (JAK1, TYK2, STAT1, and IRF9). These deficiencies are usually fatal in mice due to massive systemic spread of infection and the inability of the host immune system to control the acute infection, as demonstrated by *in vivo* studies using different transgenic mouse strains [54]. Further, IRF1 and type I IFN were shown to cooperatively control acute MHV68 infection [55], while IRF3 and IFNAR are necessary to attenuate transcription of MHV68 in primary macrophages [56].

Similar to human gammaherpesviruses, MHV68 has established a fine balance with the host immune system by using different cell types to establish and maintain latency [17,57–60]. MHV68 antagonizes the type I IFN response to promote lytic infection and establish latency, resulting in limited type I IFN secretion from *in vitro* infected cells [61,45,46]. Nevertheless, the type I IFN response plays an important role in controlling acute MHV68 infection and reactivation in mice. Dutia et al. showed that *Ifnar1^-/-^* mice are highly susceptible to MHV68 infection, while wild-type mice are able to efficiently control it [32]. Following a high dose intranasal infection (4✕10E6 PFU), around 80-90% of *Ifnar1^-/-^* mice succumbed to infection, whereas ∼ 50% of mice survived the infection at a lower dose (4✕10E3 PFU). Furthermore, the viral titers in the lungs of *Ifnar1^-/-^* mice were 100-1000 fold higher than WT, at both high and low dose infections, and MHV68 disseminated faster systemically in the absence of the IFN-I response, as indicated by higher viral titers in the spleen of *Ifnar1^-/-^*following intranasal infection.

In the present study, we similarly observed that *Ifnar1^-/-^* mice showed a significantly higher frequency and number of infected B cells compared to WT mice and the infection spread faster to the spleen in *Ifnar1^-/-^* mice, as the spleen was extensively infected on day 7 post s.c. footpad infection. The type I IFN response could contribute to infection containment by inducing the expression of various ISGs that inhibit viral replication, cell-to-cell spread, and/or virion production. Additionally, in the context of gammaherpesviruses, the infected lymphocytes could contribute to the spread of infection by migrating from the lymph nodes and entering the blood circulation. MHV68-encoded non-coding RNA, TMER4, was shown to promote the egress of infected B cells from lymph nodes, thus being necessary for the infection of circulating blood [62,63].

Although we were not able to detect the expression of *IFNα* or *IFNβ* genes in the infected lymph nodes, we could detect increased expression of *ISG15* and *MX1* in the infected lymph nodes in WT mice at 18-24 hours p.i., while the expression of these ISGs was decreased in *Ifnar1^-/-^* mice. *ISG15* is a ubiquitin-like protein that is rapidly upregulated after stimulation with type I IFN or viral infection and can be found in a conjugated form in the host or viral proteins or in an unconjugated extracellular form, acting as an immune modulator [64]. *ISG15* can limit viral progeny production by inhibiting viral enzymes, while simultaneously acting as a chemoattractant and modulator of immune cells. Lenschow *et al*. showed that *ISG15^-/-^* mice have increased susceptibility to *in vivo* MHV68 infection. Despite no change in mortality, *ISG15^-/-^* mice nevertheless demonstrated higher viral titers in the lungs and the liver on days 3 and 4 p.i.. Interestingly, they managed to clear MHV68 infection by day 6 p.i. similar to WT mice [65], indicating that ISG15 protects the host from MHV68 acute infection but its expression is not crucial for the control of the acute phase *in vivo*. Additionally, we detected higher expression of the chemokine CXCL10 in WT vs *Ifnar1^-/-^* lymph nodes. CXCL10, also known as interferon γ-induced protein 10 kDa (IP-10), is an important chemokine for the initiation of inflammation. CXCL10 is secreted by various cell types in response to IFN-γ and recruits and chemoattracts activated T cells, B cells, and NK cells via the CXCR3 chemokine receptor [66]. NK cells are one of the main producers of IFN-γ early after viral infections, and we observed that NK cells were recruited to the site of infection in *Ifnar1^-/-^* mice following MHV68 infection similarly to WT mice, which indicates that IFN-γ secreting NK cells could contribute to initiating inflammation in the absence of type I IFN signaling.

We also observed that the expression of m*OASL2* was decreased in *Ifnar1^-/-^* mice. mOASL2 is a murine homolog of human OASL (oligoadenylate synthetase-like protein) and its role in controlling MHV68 *in vivo* infection has not been studied. mOASL2 has been shown to have opposite effects following infection with DNA and RNA viruses [67]. While it inhibits RNA virus infection by enhancing the function of the RNA sensor RIG-I [68], mOALS2 inhibits type I IFN induction by binding to and inhibiting the DNA sensor cGAS. Hence, mOASL2 in the context of DNA virus infection might promote viral replication [69].

In addition to preventing viral replication, type I IFN also helps shape an effective adaptive immune response. The impact of the type I IFN response on the development of the adaptive immune response appears to be context-dependent and complex, and a better understanding may help rational vaccine design [70]. In this study, we observed that the main effector populations of the adaptive immune response, including effector CD8^+^ T cells as well as CD4^+^ Tfh cells and GC B cells expanded in the lymph nodes following MHV68 infection in both WT and *Ifnar1^-/-^* mice. This indicates that the co-stimulatory effect of the IFN response could be partially compensated by the presence of numerous infected cells.

In WT mice, CD8^+^ T cells play a crucial role in controlling different phases of MHV68 infection, including acute and latent infection as well as reactivation, via secretion of perforin and granzyme granules and IFNγ [71,41,72]. In addition to indirectly affecting T-cell priming and activation by upregulating the expression of MHCI and co-stimulatory molecules (CD80, CD86) on DCs, the type IFN response can directly promote or inhibit T cell proliferation and cytokine production [73–75]. Interestingly, Jennings *et al*. showed that type I IFN signaling is essential for the control of latent MHV68 infection and reactivation by CD8^+^ T cells as the expression of TNF-α, IFNγ, and IL2 was decreased in antigen-specific CD8^+^ T cells in *Ifnar1^-/-^* mice. Suppressing the high levels of viral reactivation did not improve CD8^+^ T cell function during latency, suggesting that IFN-I contributes to CD8^+^ T cell function indirectly by suppressing viral replication and possibly preventing T cell exhaustion in the context of MHV68 infection [33,76].

In addition to type I IFN evasion, herpesviruses have developed various strategies to evade the NK cell-mediated immune response by suppressing the signaling of activating receptors and triggering the signaling of inhibitory receptors. Nevertheless, NK cells might play an important role in controlling human and murine herpesvirus infections [38,77–79]. There are currently only a few studies on the role of NK cells in the context of MHV68 infection, and the mechanisms by which MHV68 evades the NK cell-mediated immune response are not very clear.

The MHV68-encoded K3 protein has been shown to downregulate MHC-I expression to evade antigen presentation and ultimately CTL-mediated killing of infected cells [80]. Such downregulation of MHC-I expression on infected cells could lead to their recognition by NK cells as a target. Surprisingly, studies by Usherwood et al. and Thomson *et al*. have shown that although NK cells are activated, expanded, and recruited to the site of infection following MHV68 infection in C57BL/6 mice, they do not significantly contribute to the control of acute or latent MHV68 infection [51,52]. These studies reported that NK cell depletion does not lead to significantly higher viral loads in the lungs compared to control mice following i.n. infection. Additionally, NK cells appear to be redundant for the development of the adaptive immune response, in particular for MHV68-specific CD8^+^ T cells. In another study, they proposed that primed virus-specific CD4+ T cells migrate to the lungs and activate local antigen-presenting cells (APCs) by secreting IFN-γ. The activated APCs then recruit and activate NK cells, presumably by secreting IL-12, and IL-18, and activated NK cells then contribute to the killing of infected cells and suppression of viral replication via IFN-γ secretion, suggesting a CD4^+^ T cell-NK cell axis contributing to the control of MHV68 infection in the lungs [81]. Notably, during MCMV infection in the lymph nodes, NK cells restrict lytic infection of subcapsular sinus macrophages in infected lymph nodes following s.c. footpad infection, suggesting that NK cells might contribute to the immune response in a tissue-specific manner [82]. Additionally, NK cells were already established as a second innate barrier alongside type I IFN during acute lymph node infection, showing that NK cells are recruited to MHV68-infected popliteal lymph nodes and that their depletion increases the number of infected cells [38]. This is in line with our observations showing MHV68 infection in the lymph node also being regulated by both type I IFN and NK cells. Our direct comparison of NK cell recruitment in WT and *Ifnar^-/-^* mice provides evidence that type I IFN signaling is not required for NK cell accumulation in the lymph node following MHV68 infection.

In this study, we visualized interactions between NK cells and MHV68-infected cells in infected lymph nodes using Ncr1^gfp^ reporter mice. Similar to previous studies, we observed that NK cells were recruited to infected lymph nodes early following infection. However, we have now shown that they make no detectable clusters or long-lasting contacts with MHV68-infected cells. Additionally, we did not observe NK cell-mediated contact-dependent killing in our studies. One of the suggested possible mechanisms by which MHV68-infected cells can evade NK cell-mediated contact-dependent cytotoxicity is via up-regulation of the inhibitory receptor CEACAM1 on the surface of infected cells [83].

We showed that NK cell depletion increased the number of infected cells in infected lymph nodes on day 4 p.i., suggesting that NK cells may contribute to controlling MHV68 infection in a contact-independent manner. NK cells can activate other immune cells indirectly by producing various pro-inflammatory cytokines that could clear lytic infections including but not limited to IFNγ. In addition, NK cells can perform antibody-dependent cytotoxicity [84]. NK cells recognize infected cells in which membrane-surface antigens are bound to an antigen-specific antibody via their CD16 receptors that bind to the constant part of the antibody. As the antibodies against MHV68 infection are developed at later time points p.i., it is possible for NK cells to kill MHV68 infected cells at later time points in a contact-dependent manner.

### Conclusion and future directions

In this study, we used MHV68 s.c. footpad infection in WT and transgenic mice as a model to investigate the role of the type I IFN and NK cell-mediated immune responses in controlling acute gammaherpesvirus infection in the lymph nodes. A better understanding of gammaherpesvirus control specifically in lymph nodes is required to design optimal immunotherapies of e.g. transplant-related lymphomas that are often associated with human gammaherpesviruses [85,86]. In addition, our MHV68 infection model is ideal to investigate the interactions of other immune cells, e.g. cytotoxic CD8^+^ T cells, with MHV68-infected motile B cells and their efficiency in killing these cell types in future studies. Furthermore, our model might be helpful to better understand aspects of immune dysregulation relevant to e.g. secondary EBV-triggered HLH, a very difficult to treat condition [87]. Notably, our model may help to elucidate aspects of immune dysregulation relevant to secondary EBV-triggered HLH, rather than fully recapitulating diagnostic HLH pathology.

Type IFN played a crucial role in protecting the mice from detrimental MHV68 infection by controlling the number of infected B cells as well as containing the infection and preventing extensive systemic spread of the infection, partially via induction of the ISGs. The next step would be to identify the cell types that respond to IFN in this context, as well as their ISG expression profile. The mechanisms by which MHV68-infected B cells circumvent NK cell-mediated killing needs to be further investigated. It would be interesting to study the contact-independent effect of NK cells in more detail by studying their cytokine profile and NK-mediated indirect activation of other immune cells, e.g. macrophages. Additionally, it would be relevant to study other hallmarks of HLH in our model, to test the relevance of the IFN response on full-blown HLH pathology (e.g. cytopenia, ferritin release and organ failure).

Taken together, a clear visualization of gammaherpesvirus-infected cells and the local immune cell response allows a better understanding of the early events of host colonization by a persisting viral pathogen that establishes chronic infections even in immunocompetent individuals.

## Materials and Methods

### Ethics Statement

All animal experiments were performed in accordance with the German Tierschutzgesetz and the Directive of the European Parliament and the Council on the Protection of Animals Used for Scientific Purposes (2010/63/EU). The animal studies were approved by the Niedersächsisches Landesamt für Verbraucherschutz und Lebensmittelsicherheit (LAVES) under the reference numbers 33.12-42502-04-17/2737 and 33.19-42502-04-17/2657. Mice were bred and housed at the central animal facility of Hannover Medical School under specific pathogen-free conditions. Mice were housed in individually ventilated cages with a light-dark cycle of 14:10 hours and humidity of 55% ± 5% at 22°C ± 2°C. Food and water were provided *ad libitum*.

### Animal Experiments

C57BL/6 (C57BL/6NCrl, Charles River Laboratories), *Ifnar1^-/-^* (B6.129S2-*Ifnar1^tm1Agt^*/Mmjax, Jackson Laboratory, 32045), and Ncr1^gfp^ (B6; 129-*Ncr1^tm1Oman^*/J, Jackson Laboratory, 022739) mice were used for experiments. Briefly, mice were anesthetized via intraperitoneal (i.p.) injection of a ketamine-xylazine solution according to the approved doses (50 mg ketamine/kg and 10 mg xylazine/kg of body weight). Anesthetized mice were infected with MHV68-DsRed via subcutaneous (s.c.) footpad injection with 10^6^ PFU in a 20 µl volume of sterile PBS per footpad. Mice were euthanized by CO_2_ inhalation and cervical dislocation before organ isolation.

### MHV68-DsRed reporter virus

The MHV68-DsRed reporter virus was designed by Prof. Craig Forrest (The University of Arkansas for Medical Sciences). In brief, the DsRed-Express locus, under control of the LTR promoter, was inserted between ORF57 and ORF58 in the MHV68 genome. Virus stocks were amplified in M2-10B4 cells (CRL-1972). To concentrate the virus, supernatant from infected M2-10B4 was centrifuged (3 h, 26,000 x g, 4°C). After centrifugation, the supernatant was discarded, and the pellet was soaked in virus standard buffer (50 mM Tris-HCl, pH 7.8, 12 mM KCl, 5 mM Na_2_EDTA) overnight at 4°C. The pellet was resuspended, layered on a 10% Nycodenz cushion, and centrifuged (4 h, 46,000 x g, 4°C) to purify the virus particles. After centrifugation, the pellet was resuspended in virus standard buffer. Virus stocks were titred on M2-10B4 cells, and the titer was defined using median (50%) tissue culture infective dose (TCID50) as described [88,61].

### Flow cytometry analysis

Organs were harvested and a single cell suspension was prepared. Viable cells were counted with a Vi-cell XR cell viability analyzer (Beckman) and 2x10^6^ viable cells were used per labeling reaction. Cells were transferred to 96-well V-bottom plate, resuspended in 50 µl blocking solution (PBS supplemented with 5% rat serum and 5% Fc block) and incubated for 10-15 minutes on ice. Next, the cells were protected from light and labeled with a multicolor antibody mix including ZOMBIE NIR Viability dye (Biolegend), CD4-BUV737 (BD-RM4-5), CD138-BV421 (Biolegend, 281-2), CD3e-Pacific Blue (Biolegend, 17A2), IgD-BV510 (Biolegend, 11-26c.2a), CD11b-BV570 (Biolegend, M1/70), CD27-BV650 (Biolegend, LG.3A10), CD62L-BV711 (Biolegend, MEL-14), NK1.1-BV785 (Biolegend, PK136), CD44-FITC (Biolegend, IM7), CD95-PE-Cy7 (BD, Jo2), CXCR5-PE (Invitrogen, SPRCL5), CD19-Spark Blue 550 (Biolegend, 6D5), PD-1-BB700 (BD, RMP1-30), GL7-PerCP/Cy5.5 (Biolegend), CD11c-APC (Biolegend, N418), CD8a-Alexa Fluor 647 (Biolegend, 53-6.7) for 30 minutes. The cells were then washed and analyzed with Cytek Aurora Spectral Flow cytometer equipped with 5 lasers (355 nm, 405 nm, 488 nm, 561 nm, and 640 nm). Data were analyzed with FCS Express 7 and statistical analysis was performed with Graphpad Prism 8.

### NK cell depletion

NK cells were depleted by i.p. injection of 250 µg of anti-NK1.1 antibody (clone PK 136) in 100 µl of sterile PBS 24 hrs prior to s.c. footpad infection with MHV68-DsRed and 0, 2 and 4 days post-infection. NK cell depletion was confirmed in each experiment by flow cytometry of spleen and peripheral lymph nodes from the NK-depleted and control groups.

### Two-photon microscopy

Lymph nodes were excised and fixed in a customized tissue chamber using tissue adhesive [89,26]. The chamber was then superfused with oxygenated (95% O_2_/5% CO_2_) RPMI 1640 medium supplemented with 4 g/L glucose and placed on a 37°C hotplate. Imaging was performed using a TriM Scope (LaVision Biotec) equipped with an upright microscope (BX51; Olympus) and 20X and 10X water immersion objectives. Imaging was performed using Mai Tai and InSight tunable lasers (Spectra-Physics). The wavelengths were set to 1125 nm for the Insight laser and 910 nm for the Maitai laser. Data were analyzed with Imaris 9.5.

### Immunohistochemistry analysis

Excised lymph nodes and spleens were fixed in 2% paraformaldehyde (PFA) containing 30% sucrose for 1 hour (lymph nodes) or overnight (spleen). The tissue was then embedded in a mold filled with Tissue Tek OCT (SAKURA), frozen on dry ice, and stored at -20°C. Eight μm thick cryosections were prepared and stored at -20°C. After rehydration for 5 min in TBS-T (Tris-buffered saline supplemented with Tween 20), slides were blocked with TBS-T containing 10% rat serum for 15 min, then labeled with CD169-AF488 (Biolegend, 3D6.112), CD11b-eFluor 660 (eBioscience, M1/70), B220-Cy5 (homemade), and TCRβ-PECy7 (Biolegend, H57-597) antibodies for 30 minutes at room temperature and protected from light. Afterward, slides were washed and stained with DAPI for 2 min. Slides were washed three times, mounted with Mowiol, and dried overnight in the dark at RT. Images were acquired using AxioScan Z1 (Zeiss) or AxioVert 200M (Zeiss) microscopes. Images were processed with Zen 3.1 and further analyzed using Imaris 9.5 software.

### RNA Extraction and RT-qPCR

Lymph nodes were excised and a single cell suspension was prepared by enzymatic digestion in RPMI medium supplemented with FBS (5%), HEPES (2.5 mM), DNase I (Roche, 25 µg/ml), and Collagenase D (Roche, 0.5 mg/ml). Briefly, the lymph nodes were placed in the digestion medium, disrupted using forceps, and then incubated at 37°C for 30 minutes. The reaction was stopped by adding EDTA (0.5 M). Total RNA was then isolated using the RNeasy Mini Kit Plus (Qiagen) according to the manufacturer’s instructions. The total RNA concentration and purity was measured with a NanoDrop spectrophotometer and cDNA was prepared using the SuperScript III Reverse Transcriptase kit (Invitrogen) following the manufacturer’s protocol. Quantitative PCR was conducted with the TB Green Premix Ex Taq II kit (Takara). The primers for the *ISG15, IFNα, IFNβ, MX1* and *CXCL10* genes were already described (Mboko et al., 2012; Moll et al., 2016; Stifter et al., 2019). The following primers were used for the *mOASL2* gene: For: CCAAAACGAGGTCGTCAGGAAC, Rev: AGCCACCTGTTCCCATCCCTTT and for *GAPDH*: For: CTCCACTTTGCCACTGCAAA, Rev: GTGCCAGCCTCGTCCCGTA. A StepOnePlus real-time PCR system (Applied Biosystems) was used to acquire the data. The data was analyzed with the StepOne v2.3 software.

### Statistical analysis

Statistical analysis was performed with GraphPad Prism 8. Data interpretation in this study relies primarily on effect sizes. The statistical significance of differences between groups was analyzed using the Mann-Whitney U test (p-value <0.05 was considered significant). All data were recorded in at least two independent experiments, and only pooled data are shown. No outliers were excluded. Data interpretation is primarily based on effect size estimation. Where needed and applicable, Cohen’s d is reported in the Fig legends to quantify the magnitude of observed differences.

## Acknowledgments

We thank Thomas Schulz and the Collaborative Research Centre CRC 900 team for their consistent support and a collaborative environment.

## Supporting information

S1 Movie 1: **Two-photon microscopy of MHV68-DsRed-infected cells in lymph nodes.** Wildtype mice were infected subcutaneously (s.c.) in the footpad with an MHV68-DsRed reporter virus (day 0) and sacrificed on day 7 post-infection for extraction of the lymph nodes followed by two-photon imaging. Movie 1 shows MHV68-DsRed-infected (red) and noninfected cells at 7 days post-infection in the lymph node. Movie 1 is shown from top view **(A),** lateral view **(B),** and top view without blue second-harmonic generation signal **(C)**. Scale bar: 50 µm.

S2 Movie 2: **Two-photon microscopy of MHV68-DsRed-infected cells in lymph nodes of Ncr1^gfp^ reporter mice.** Ncr1^gfp^ reporter mice, allowing fluorescent visualization of NK cells, were infected s.c. with MHV68-DsRed in the footpad. The popliteal LNs were isolated on day 3 and 5 p.i. and analyzed by two-photon microscopy. **(A)** Movie showing NK cells (green) in the MHV68-DsRed (red) infected LNs at day 3 p.i. Scale bar: 50 μm. Second-harmonic generation SHG indicates collagen (blue). **(B)** Movie showing the motility and interactions between NK cells (green) and motile MHV68-DsRed infected cells (red). Scale bar: 30 μm.

S3 Dataset: Minimal dataset generated in this study.

## Notes

### Competing Interest Statement

The authors have declared no competing interest.

